# Intradermal-delivered DNA vaccine provides anamnestic protection in a rhesus macaque SARS-CoV-2 challenge model

**DOI:** 10.1101/2020.07.28.225649

**Authors:** Ami Patel, Jewell Walters, Emma L. Reuschel, Katherine Schultheis, Elizabeth Parzych, Ebony N. Gary, Igor Maricic, Mansi Purwar, Zeena Eblimit, Susanne N. Walker, Diana Guimet, Pratik Bhojnagarwala, Arthur Doan, Ziyang Xu, Dustin Elwood, Sophia M. Reeder, Laurent Pessaint, Kevin Y. Kim, Anthony Cook, Neethu Chokkalingam, Brad Finneyfrock, Edgar Tello-Ruiz, Alan Dodson, Jihae Choi, Alison Generotti, John Harrison, Nicholas J. Tursi, Viviane M. Andrade, Yaya Dia, Faraz I. Zaidi, Hanne Andersen, Mark G. Lewis, Kar Muthumani, J Joseph Kim, Daniel W. Kulp, Laurent M. Humeau, Stephanie Ramos, Trevor R.F. Smith, David B. Weiner, Kate E. Broderick

## Abstract

Coronavirus disease 2019 (COVID-19), caused by the SARS-CoV-2 virus, has had a dramatic global impact on public health, social, and economic infrastructures. Here, we assess immunogenicity and anamnestic protective efficacy in rhesus macaques of the intradermal (ID)-delivered SARS-CoV-2 spike DNA vaccine, INO-4800. INO-4800 is an ID-delivered DNA vaccine currently being evaluated in clinical trials. Vaccination with INO-4800 induced T cell responses and neutralizing antibody responses against both the D614 and G614 SARS-CoV-2 spike proteins. Several months after vaccination, animals were challenged with SARS-CoV-2 resulting in rapid recall of anti-SARS-CoV-2 spike protein T and B cell responses. These responses were associated with lower viral loads in the lung and with faster nasal clearance of virus. These studies support the immune impact of INO-4800 for inducing both humoral and cellular arms of the adaptive immune system which are likely important for providing durable protection against COVID-19 disease.

## Introduction

COVID-19 was declared a global pandemic on March 11, 2020 by the World Health Organization. There are >15.5 million confirmed cases worldwide with the total number of deaths estimated to be >600,000 (July 24, 2020, gisaid.org). COVID-19 presents as a respiratory illness, with mild-to-moderate symptoms in many cases (~80%) (Chen et al., 2020; Huang et al., 2020). These symptoms include headache, cough, fever, fatigue, difficulty breathing, and possible loss of taste and smell. The factors involved with progression to severe COVID-19 disease in 20% of cases are unclear; yet, severe disease is characterized by development of a hyperinflammatory response, followed by development of acute respiratory distress syndrome (ARDS), potentially leading to mechanical ventilation, kidney failure, and death (Huang et al., 2020; Xu et al., 2020). The rapid development of vaccine countermeasures remains a high priority for this infection, with multiple candidates having entered the clinic in record time (Thanh Le et al., 2020).

SARS-CoV-2 is a positive-sense single stranded RNA virus belonging to the family *Coronaviridae*, genus *Betacoronavirus*, the same family as severe acute respiratory syndrome coronavirus (SARS-CoV) and Middle East respiratory syndrome coronavirus (MERS-CoV). SARS-CoV-2 is homologous to SARS-CoV, sharing around 70-80% of its genome (Lu et al., 2020). Structural components of SARS-CoV-2 include the envelope (E) protein, membrane (M) protein, nucleocapsid (N) protein, and surface spike (S) protein, which is a major immunogenic target for humoral and cellular immune responses.

Virus entry into host cells is mediated through S protein receptor binding domain (RBD) interaction with the host cell receptor angiotensin converting enzyme 2 (ACE2) and priming by the serine protease TMPRSS2 (Hoffmann et al., 2020). Recent preprints and published studies also implicate S1 binding to neuropilin 1 (NLP-1) and have identified neutralizing epitopes outside RBD (Chi et al., 2020).

SARS-CoV-2 is the third coronavirus outbreak this century. Prior work with the related coronaviruses, SARS-CoV and MERS-CoV, delineated that the Spike protein of these viruses was an important target for development of neutralizing antibodies, and in animal viral challenges vaccine targeted immunity (reviewed in (Du et al., 2009; Roper and Rehm, 2009; Thanh Le et al., 2020) (Liu et al., 2018; Muthumani et al., 2015; van Doremalen et al., 2020a).

Recently studies have described initial data for vaccines developed from inactivated viruses (Gao et al., 2020), recombinant adenoviral vectors that express the spike antigen (van Doremalen et al., 2020b), or intramuscular (IM) delivered DNA vaccine candidates (Yu et al., 2020) in non-human primates (NHPs). These studies focused on disease protection in acute models with challenge weeks after the final vaccination. Two in particular showed the ability of vaccination to lower viral load in the lungs (Gao et al., 2020; van Doremalen et al., 2020b), and one reported impact in the lungs and a trend to impact viral loads in the nose (Yu et al., 2020). While there are significant questions arising regarding the length of protective immunity from SARS-CoV2 infection, as well as the long-term protection of vaccination, there have been no studies yet reported of the ability of vaccine memory to impact challenge outcome. Here we report on the impact of an ID-delivered SARS-CoV-2 DNA vaccine in a non-human primate model where the animals were challenged 3 months post-vaccination.

Recent studies support an important role for both cellular and humoral immunity in protection against COVID-19 disease. T cell-mediated immunity has been shown to be important for other beta-coronaviruses, providing both direct protection and help in the generation of effective humoral responses (Channappanavar et al., 2014b; Gretebeck and Subbarao, 2015). Several SARS-CoV-2 studies have now reported that up to 40% of persons can exhibit mild disease which may be associated with T cell immunity (Payne et al., 2020) or humoral responses (Robbiani et al., 2020). This echoes older studies which found that neutralizing antibody levels and memory B cell responses have not been sustained in SARS survivors, while responsive T cell populations have proved more durable (Le Bert et al., 2020; Tang et al., 2011; Yang et al., 2006). Studies have indicated the levels of SARS-CoV-2 antibodies wane rapidly in convalescent subjects (Long et al., 2020).

We recently described the immunogenicity of a SARS-CoV-2 DNA vaccine (INO-4800), encoding a synthetic spike immunogen in small animal models (Smith et al., 2020). This vaccine candidate induced neutralizing and ACE2-blocking antibodies, as well as T cell responses in mice and guinea pigs. Here, we assess vaccine-induced memory T and B cell responses during the acute expansion and memory phases in rhesus macaques, as well the ability of vaccine-induced memory to impact infection in an NHP viral installation challenge following vaccination by the minimally invasive (ID-EP) route. Additionally, we addressed the growing concerns regarding the emergence of a dominant SARS-CoV2 variant G614, which now comprises more than 80% of circulating global viral strains, and is reportedly associated with increased infectivity and spread (Korber B et al., 2020). There have not yet been descriptions of vaccine induced responses driving immunity against G614 variants. We report vaccine-induced humoral and cellular immunity, including neutralizing antibody responses against both the SARS-CoV-2 D614 virus as well as the new G614 strain. Upon SARS-CoV-2 challenge more than 3 months following final-dose, vaccinated macaques exhibited a rapid recall response against multiple regions of the S protein. This anamnestic response was characterized by expansion of neutralizing antibody responses, including those against the now dominantly circulating G614 variant, as well as the rapid expansion of T cell responses. The immune responses induced by INO-4800 were associated with disease protection.

## Results

### Induction of memory humoral and cellular immune responses in INO-4800 immunized non-human primates

NHPs are a valuable model in the development of COVID-19 vaccines and therapeutics as they can be infected with wild-type SARS-CoV-2, and present with early infection that mimics aspects of human disease (Chandrashekar et al., 2020; Gao et al., 2020; Yu et al., 2020). Rhesus macaques (n=5) received two immunizations of INO-4800 (1 mg), at Week 0 and Week 4 (**Figure 1A**). Naïve control animals (n=5) did not receive vaccine. Humoral and cellular immune responses were monitored for 15 weeks (~4 months) following prime immunization for memory responses. All animals seroconverted following a single INO-4800 immunization, with serum IgG titers detected against the full-length S1+S2 extracellular domain (ECD), S1, S2, and RBD regions of the SARS-CoV-2 S protein (**Figure 1B and C**). Cross-reactive antibodies were also detected against SARS-CoV S1 protein and RBD, but not MERS-CoV (**Supplementary Figure 1**). SARS-CoV-2-reactive IgG against the ECD and RBD were detected in bronchoalveolar lavage (BAL) washes at Week 8, 4-weeks following the 2nd immunization dose (**Supplementary Figure 2**).

**Figure 1.**
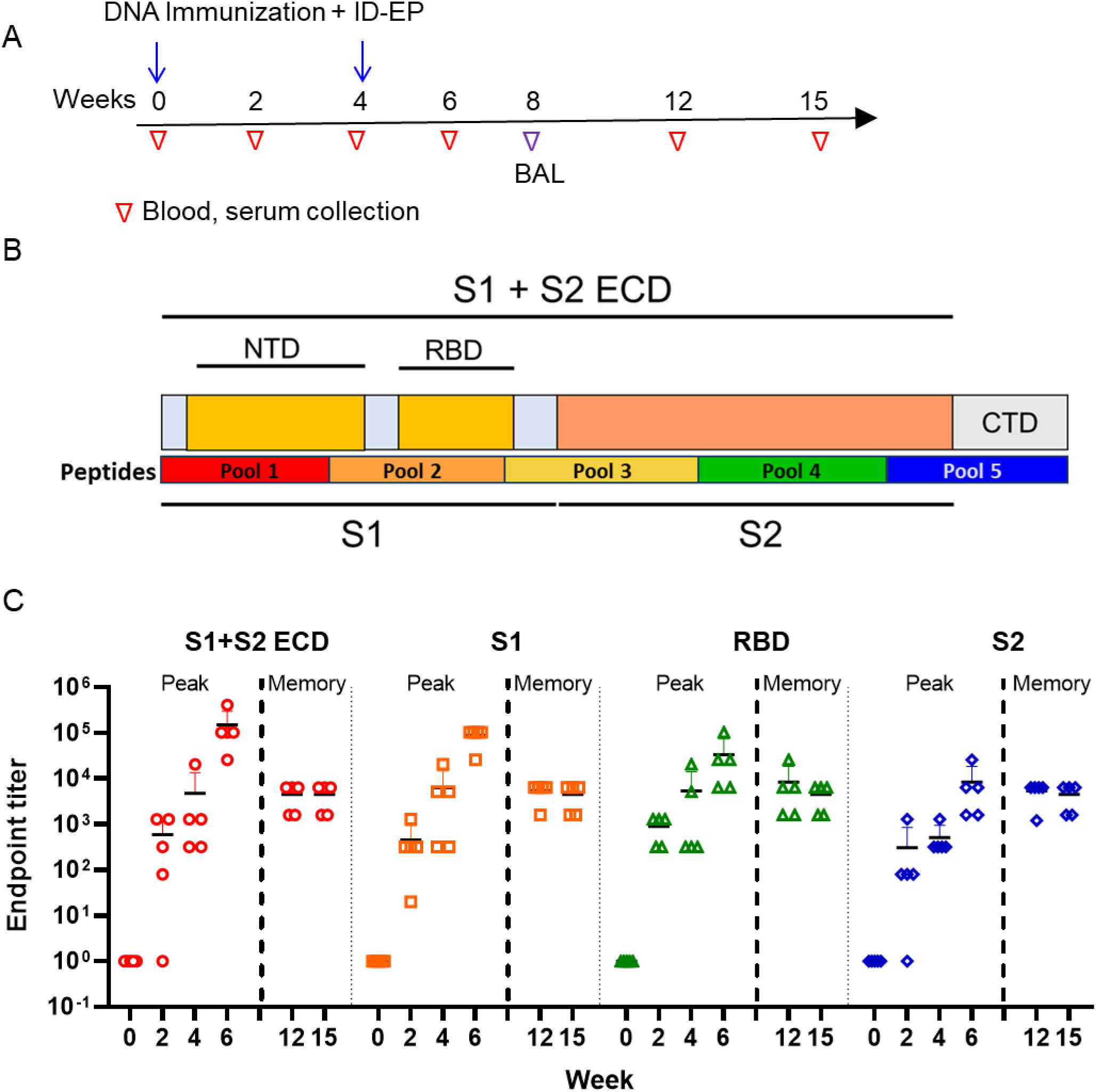
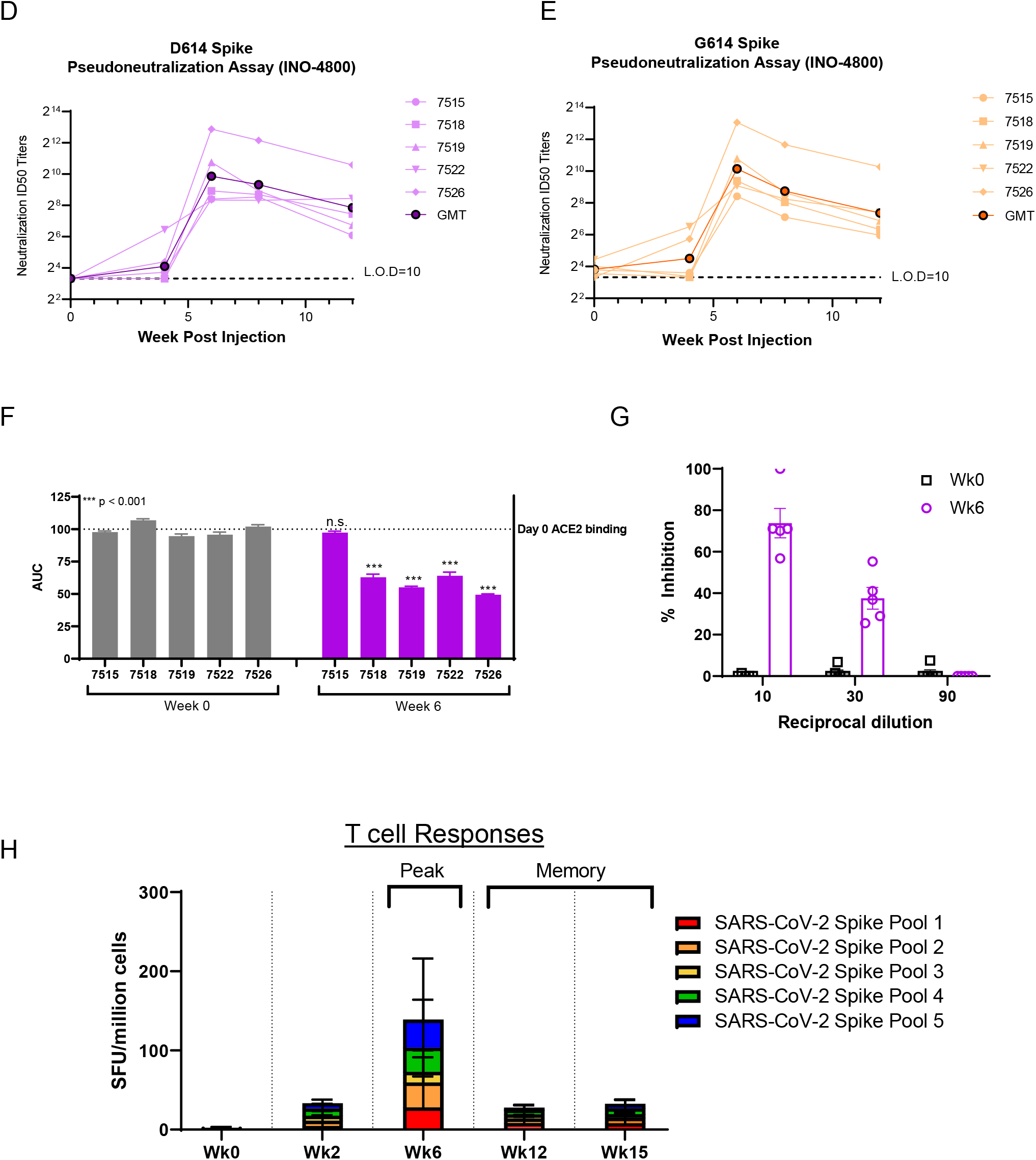
Humoral and cellular immune responses in rhesus macaques. (**A**) The study outline showing the vaccination regimen and sample collection timepoints. (**B**) Schematic of SARS-CoV-2 spike protein. (**C**) SARS-CoV-2 S1+S2 ECD, S1, RBD and S2 protein antigen binding of IgG in serially diluted NHP sera collected. Data represents the mean endpoint titers for each individual NHP. (D&E) Pseudoneutralization assay using NHP sera, showing the presence of SARS-CoV-2 specific neutralizing antibodies against the D614 (**D**) and G614 (**E**) variants of SARS-CoV-2. (**F&G**) Serum collected at Week 6 from INO-4800 vaccinated NHPs inhibited ACE2 binding to SARS-CoV-2 Spike protein. A plate-based ACE2 competition assay (**F**) and a flow-based ACE2 competition assay (**G**) showing inhibition of ACE2 binding by NHP sera. (**H**) T cell responses were measured by IFN-γ ELISpot in PBMCs stimulated for 20h with overlapping peptide pools spanning the SARS-CoV-2 Spike protein. Bars represent the mean + SD.

SARS-CoV-2 pseudovirus-neutralizing antibodies were detected in the serum of vaccinated animals at week 12 following immunization (**Figure 1D**). These memory titers were comparable to those observed in other reported protection studies in macaques performed at the acute phase of the vaccine-induced immune response (Gao et al., 2020; van Doremalen et al., 2020b; Yu et al., 2020) and those reported in the sera of convalescent patients (Ni et al., 2020; Robbiani et al., 2020). During the course of the COVID-19 pandemic, a D614G SARS-CoV-2 spike variant has emerged that has potentially greater infectivity, and now accounts for >80% of new isolates (Korber B et al., 2020). There is a concern that vaccines developed prior to this variant’s appearance may not neutralize the D614G virus. There is also concern that persons infected with earlier strains, may not have immunity against the G614 variant. Therefore, we evaluated neutralization against this new variant using a modified pseudovirus developed to express the G614 spike protein (**Figure 1E**). Similar neutralization ID50 titers were observed against both D614 and G614 spikes, supporting induction of functional antibody responses by the vaccine against both SARS-CoV-2 virus variants.

To investigate specific receptor-blocking neutralizing antibody activity, we assessed the capacity to occlude ACE2 by vaccine-induced sera. We recently developed a receptor-blocking assay and demonstrated that it correlates with pseudovirus neutralization titers (*Walker et al. accepted*). In this assay, anti-spike antibodies are bound to SARS-CoV-2 spike protein in an ELISA plate followed by addition of a soluble version of the ACE2 receptor. Sera containing antibodies which can obstruct the receptor binding site on the spike protein will have lower Area Under the Curve (AUC) values in our assay. We observe that sera from 80% of immunized NHPs had reduced AUC indicating the presence of antibodies that can block the SARS-CoV-2 spike protein from interacting with ACE2 (**Figure 1F**). An independent flow cytometry-based assay was also performed to further study the Spike-ACE2 interaction. ACE2-expressing 293T cells were co-incubated with Spike with or without the presence of sera. Spike binding to ACE2 was detected by flow cytometry. In this assay, 100% of macaques inhibited the Spike-ACE2 interaction, with 53-96% inhibition of the spike-ACE2 interaction at a 1:10 dilution and 24-53% inhibition at a 1:30 dilution (**Figure 1G**).

INO-4800 immunization induced SARS-CoV-2 S antigen reactive T cell responses against all 5 peptide pools with T cells responses peaking at Week 6, two weeks following the second immunization (0-518 SFU/million cells, with 4/5 NHPs showing positive responses) (**Figure 1H**). Distinct immunogenic epitope responses were detected against the RBD and S2 regions (**Figure 1B and C**). Interestingly, cross-reactive T cells responses were also detected against the SARS-CoV spike protein (**Supplementary Figure 3A**). However, we did not observe cross-reactivity to MERS-CoV spike peptides, which is consistent with the lower sequence homology between SARS-CoV-2 and MERS-CoV (**Supplementary Figure 3B**).

### Vaccine induced immune recall responses upon SARS-CoV-2 challenge in non-human primates

INO-4800 immunized macaques and unvaccinated controls were challenged with SARS-CoV-2 13 weeks (~3 months) post-final immunization (Study Week 17, **Figure 2A**). NHPs received a challenge dose of 1.1×10^4^ PFU of SARS-CoV-2 Isolate USA-WA1/2020 by intranasal and intratracheal inoculation as previously described (Chandrashekar et al., 2020; Yu et al., 2020). One day after viral challenge, 3/5 of INO-4800 vaccinated animals displayed an increase in antibody titers against the SARS-CoV-2 full-length ECD. By day 7, all vaccinated animals had an increase in antibody titers against the full length ECD, S1, RBD, and S2 proteins (**Figure 2B**). Seven days post-challenge, robust geometric mean endpoint titers for the ECD ranging from 409,600 - 1,638,400 were observed in immunized animals, compared with the naïve group which displayed seroconversion of only 1/5 animals (GMT 100) (**Figure 2B**). Antibody responses continued to increase against SARS-CoV-2 proteins 14 days post-challenge. A significant increase in pseudoneutralization titers (>20-fold increase) was observed in all INO-4800 immunized animals against both D614 and G614 spike variants by day 7 post-challenge, compared to unimmunized animals which only developed low neutralizing titers (**Figure 2C**).

**Figure 2.**
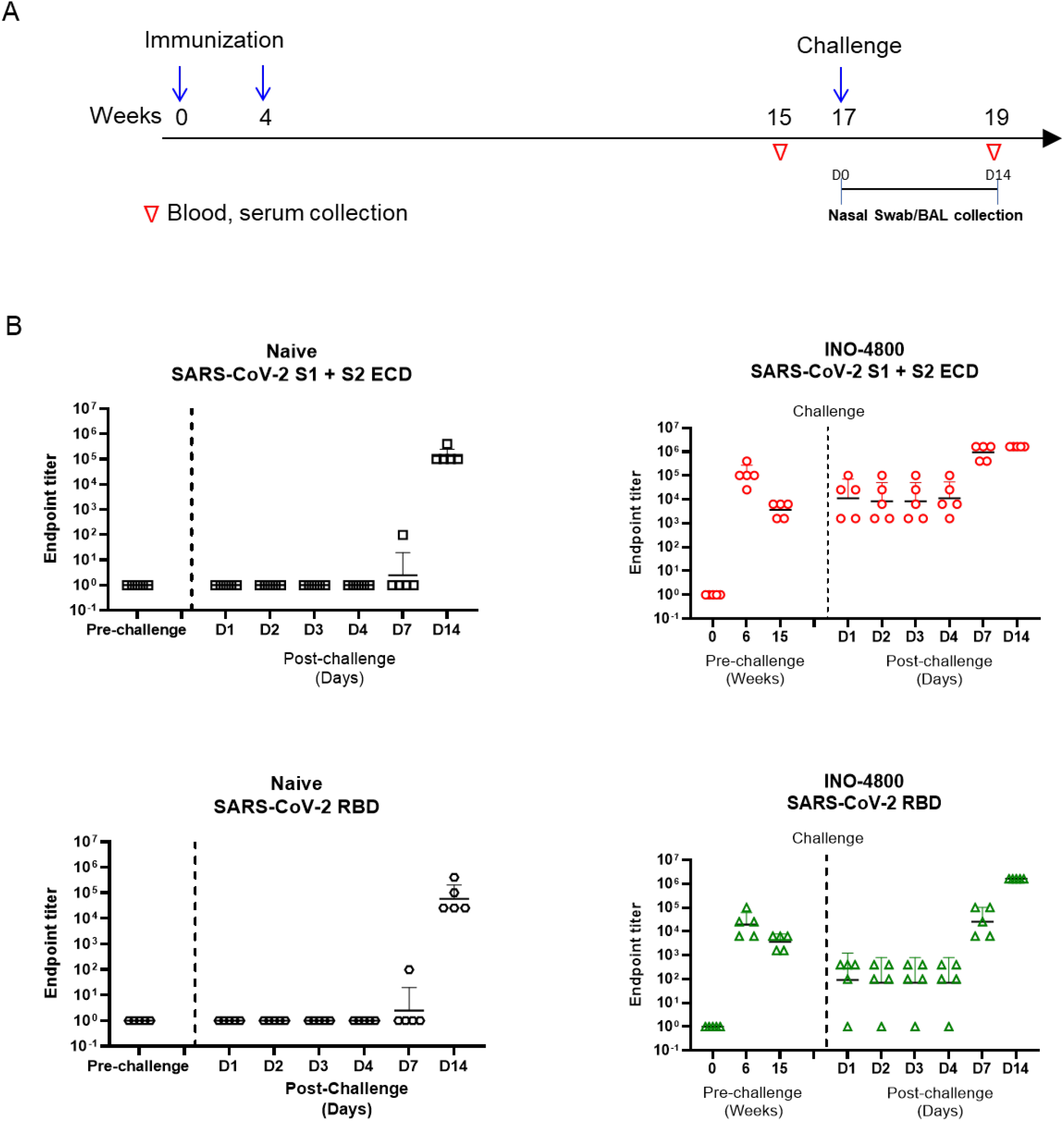
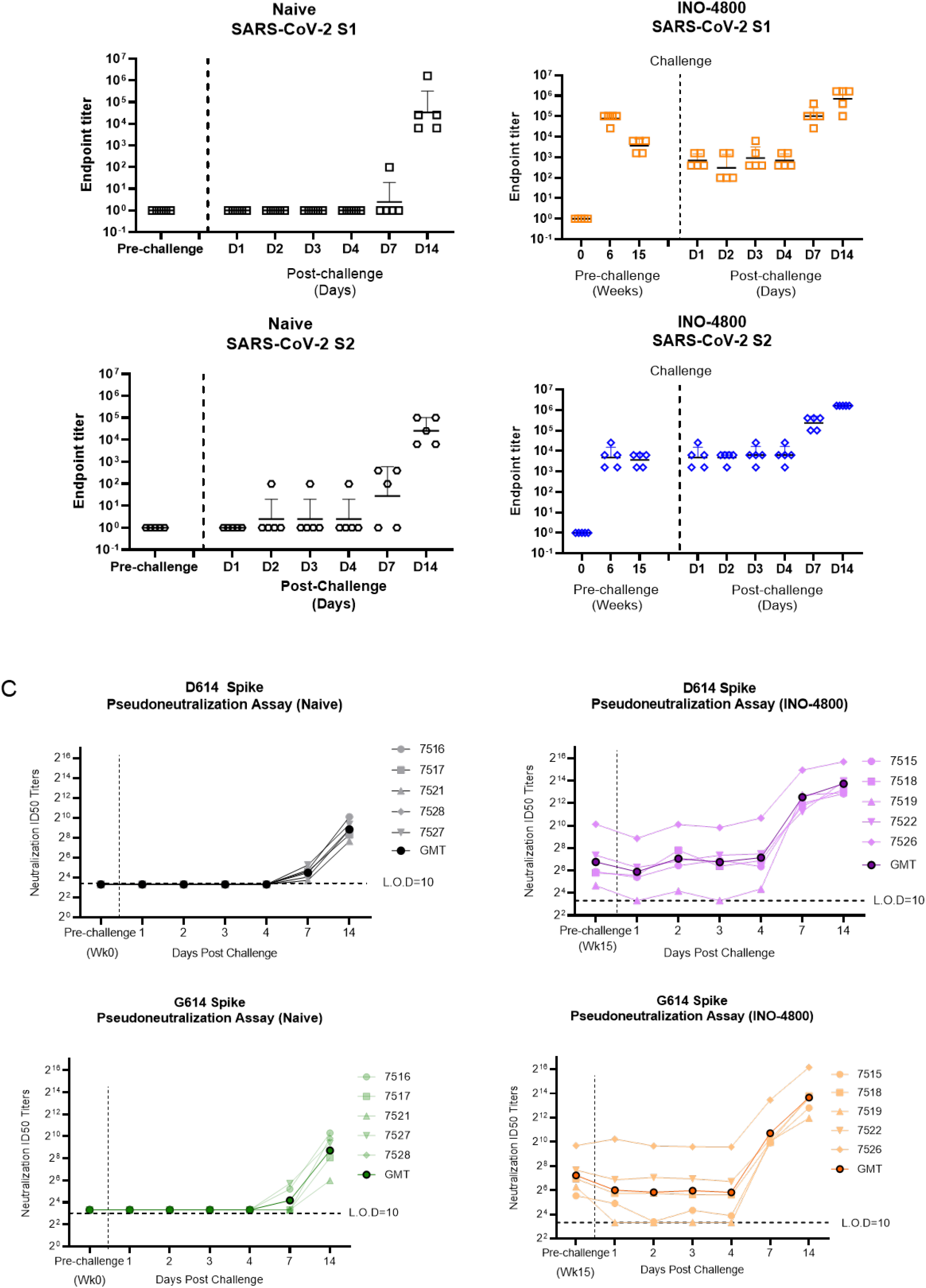
Recall of humoral immune responses after viral challenge. (**A**) Study outline. (**B**) IgG binding ELISA. SARS-CoV-2 S1+S2 and SARS-CoV-2 RBD protein antigen binding of IgG in diluted NHP sera collected prior to challenge and post challenge in INO-4800 vaccinated (right panels) and naïve animals (left panels). (**C**) Pseudo-neutralization assay showing the presence of SARS-CoV-2 specific neutralizing antibodies against the D614 and G614 variants of SARS-CoV-2 before and after viral challenge in unvaccinated (left panels) and INO-4800 vaccinated (right panels).

Cellular responses were evaluated before and after challenge. At week 15, IFN-γ ELISpot responses had contracted to memory but persisted (**Figure 1H**). Following challenge, T cell responses increased for all animals in the vaccinated group (~218 SFU/million cells average) supporting rapid recall expansion of immunological T cell memory (**Figure 3 and Supplementary Figure 4**).

**Figure 3.**
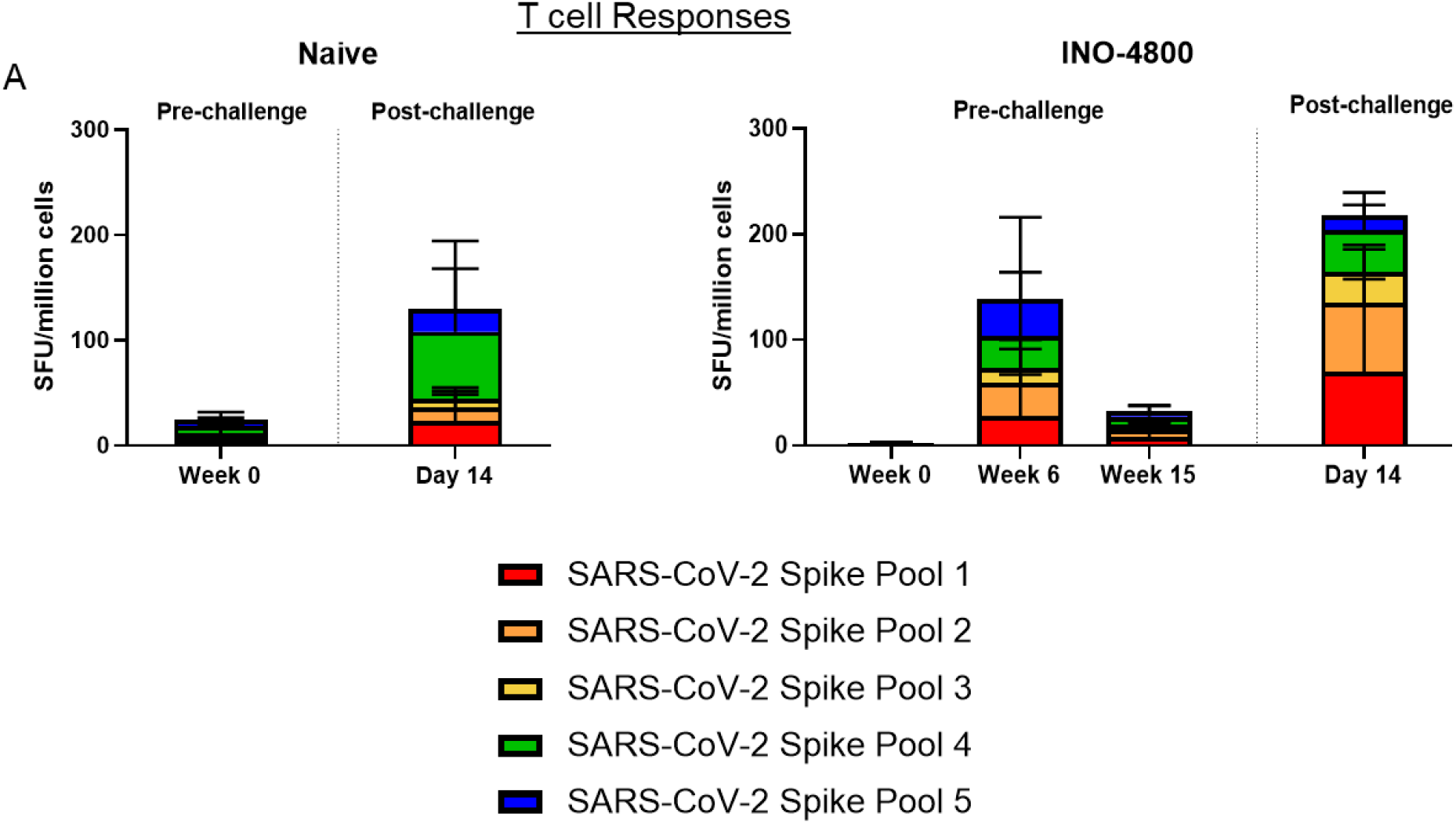
Recall of cellular immune responses after viral challenge. (**A**) T cell responses were analyzed pre and post challenge with SARS-CoV-2 virus by IFNγ ELISpot in PBMCs stimulated with overlapping peptide pools spanning the SARS-CoV-2 spike protein. Bars represent the mean + SD. Left panel naïve animals, right panel INO-4800 vaccinated animals.

### Protective efficacy following SARS-CoV-2 challenge

At earlier time points post challenge, viral mRNA detection does not discriminate between input challenge inoculum and active infection, while subgenomic (sgmRNA) levels are more likely representative of active cellular SARS-CoV-2 virion replication (Wolfel et al., 2020; Yu et al., 2020). As such, SARS-CoV-2 sgmRNA was measured in unvaccinated control and INO-4800 vaccinated macaques following challenge (**Figure 4**). Peak viral sgmRNA loads in the BAL were significantly lower in the INO-4800 vaccinated group **(Figure 4A and B)**, along with significantly lower viral RNA loads at day 7 post-challenge (**Figure 4C**), indicating protection from lower respiratory disease. While sgmRNA was detected in the nasal swabs of both the control and INO-4800 vaccinated animals **(Figure 4D-F)**, viral mRNA levels trended downwards in INO-4800 vaccinated animals by more than 2 logs (**Figure 4F**) and were achieved sooner on average. Overall, the reduced viral loads following exposure to SARS-CoV-2 infection at 17 weeks after immunization show an important durable impact mediated by the vaccine.

**Figure 4.**
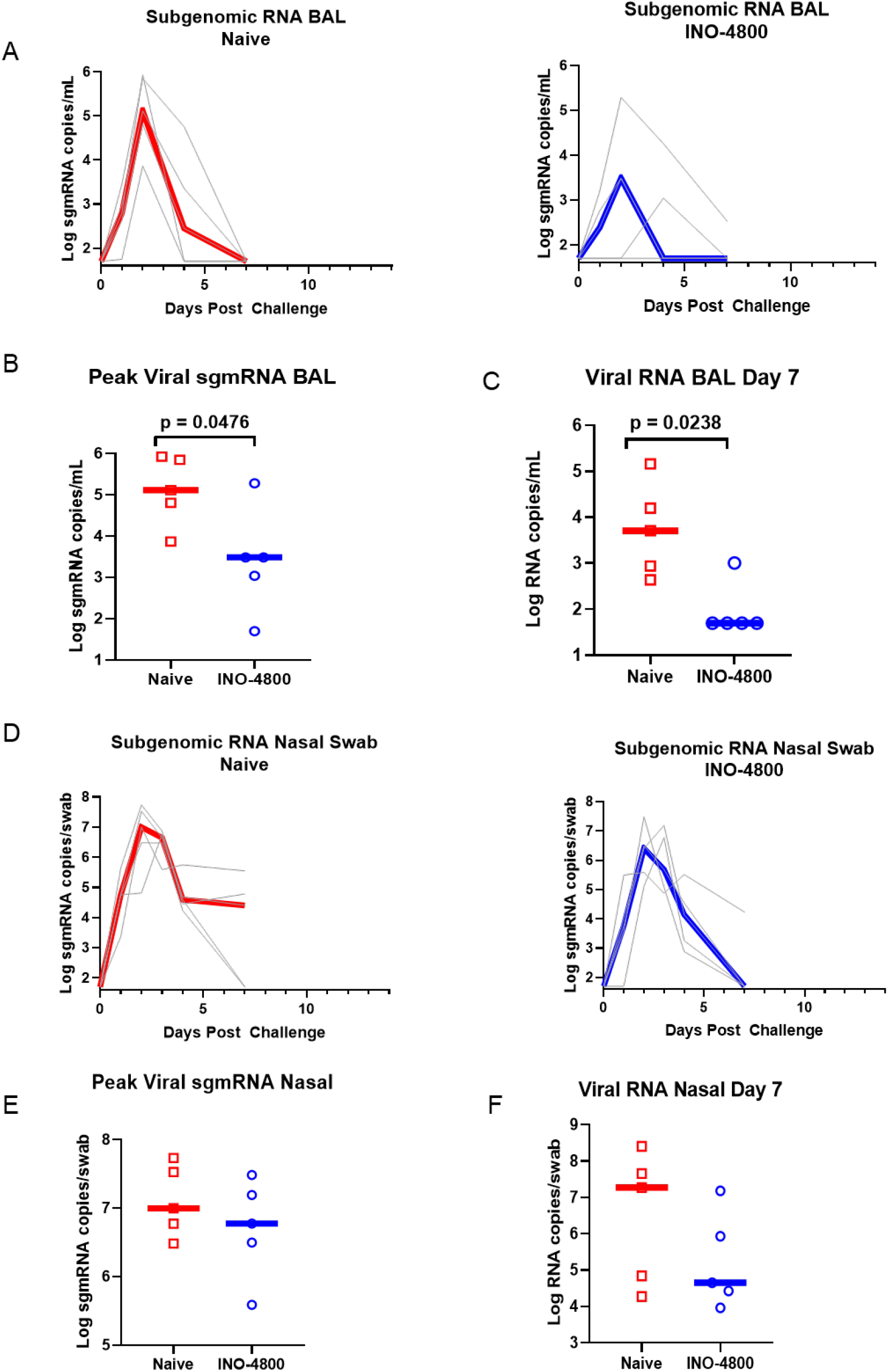
Viral loads in the BAL fluid and Nasal swabs after viral challenge. At week 17 naïve and INO-4800 immunized (5 per group) rhesus macaques were challenged by intranasal and intratracheal administration of 1.1 x 10^4^ PFU SARS-CoV-2 (US-WA1 isolate). (**A**) Log sgmRNA copies/ml in BAL in naïve (left panel) and INO-4800 vaccinated animals (right panel). (**B**) Peak sgmRNA and (**C**) viral RNA in BAL 7 days post-challenge. (**D**) Log sgmRNA copies/ml in nasal swabs in naïve (left panel) and INO-4800 vaccinated animals (right panel). (**E**) Peak sgmRNA and (**F**) viral RNA in nasal swabs 7 days post-challenge. Red and blue line represent median values.

## Discussion

The COVID-19 pandemic continues to devastate global health creating a destabilizing environment. Although therapies are progressing through multiple stages of development, the need for preventative vaccine approaches for SARS-CoV-2 remains significant. Importantly, multiple vaccines candidates are being advanced to the clinic as potential options for protection against infection and or disease (reviewed in (Funk et al., 2020; Thanh Le et al., 2020)). Recently acute challenge studies in NHPs receiving different experimental vaccines have been reported (Gao et al., 2020; van Doremalen et al., 2020b; Yu et al., 2020). These studies describe reduction in viral loads in the lower airways, and a trend to lower viral loads in the nose by an IM DNA delivered full length spike Ag. However, protective efficacy at time points following contraction of the acute phase immune response and establishment of immune memory have not been previously studied. This is also true looking at prior NHP challenge in the MERS or SARS models (Muthumani et al., 2015; Roberts et al., 2008; van Doremalen et al., 2020a). Accordingly, understanding the impact of vaccine-induced memory on challenge is of importance.

In the current study, we evaluated the durability of an ID-delivered full-length Spike antigen vaccine (INO-4800) in rhesus macaques. The vaccine induced broad Spike binding antibodies to multiple forms of the antigen, as well as rapid induction of potent neutralizing antibodies, and cellular immune responses. Macaques additionally elicited cross-reactive humoral and cellular responses to the SARS-CoV S protein. We further demonstrated protective efficacy more than 3 months post-final immunization, observing establishment of anamnestic immune responses and reduced viral loads upon challenge in vaccinated macaques. Recently, Yu et al. evaluated a full length Spike Ag DNA vaccine candidate, administered by 2 x 5 mg IM doses, in NHPs (Yu et al., 2020) followed by challenge at 3 weeks following the second immunization. They observed decreases in BAL viral loads and a trend to lower nasal swab subgenomic viral loads in some groups, including a group containing a full-length Spike Ag construct that shares similarity to the construct studied here using a 2 x 1 mg dose by ID administration. In a recent clinical study, we reported that ID delivery of an HIV vaccine prototype was dose-sparing and demonstrated increased tolerability and improved immune potency compared to IM DNA delivery (De Rosa et al., 2020). While limited conclusions can be drawn from such comparisons, more study in this context is warranted.

In this study, we observed an impact on induction of durable immunity and protective efficacy at a memory time point, 13 weeks post-final immunization. Importantly, reduced viral subgenomic RNA loads in the lower lung were observed. A trend of lower viral mRNA loads was also observed in the nose with more rapid clearance induced in the INO-4800 vaccinated animals. These data support that immunization and memory immunity from such a vaccine in humans might protect from severe disease in the lung as well as limit the duration of viral shedding in the nasal cavity, thus possibly lowering viral transmission.

The initial viral loads detected in control unvaccinated animals in this study were approximately 2 logs higher (10^9^ PFU/swab in 4/5 NHPs on day 1 post-challenge) than in similar published studies performed under identical conditions (~10^7^ PFU/swab) (Yu et al., 2020). Two of the prior reported NHP studies included intranasal delivery as an inoculation route for challenge (van Doremalen et al., 2020b; Yu et al., 2020). High-dose challenge inoculums are frequently employed to ensure take of infection, however such high dose challenge may artificially reduce the impact of potentially protective vaccines and interventions (Durudas et al., 2011; Innis et al., 2019). Despite these limitations, this study demonstrated significant reduction in peak BAL sgmRNA and overall viral mRNA. Wolfel et al reported nasal titers in patients of an average 6.5 x 10^5^ copies/swab days 1-5 following onset of symptoms (Wolfel et al., 2020). These titers are significantly lower than our NHP challenge dose and support the potential for the vaccine to exhibit great impact in the field against natural SARS-CoV-2 challenge.

T cell immunity has been shown to be an important mediator of protection against betacoronaviruses, and recent studies have specifically identified a role in protection against COVID-19 disease (Channappanavar et al., 2014a; Channappanavar et al., 2014b; Sekine et al., 2020; Sun et al., 2020). Sekine et al reported SARS-CoV-2-reactive T cells in asymptomatic and mild COVID-19 convalescent patients who were antibody seronegative (Sekine et al., 2020). These results from convalescent patient studies collectively suggest that vaccine candidates which could generate balanced humoral and cellular immune responses could be important in providing protection against COVID-19 disease. Our study and other published reports show that DNA vaccination with candidates targeting the full-length SARS-CoV-2 spike protein likely increase the availability of T cell immunodominant epitopes leading to a broader and more potent immune response, compared to partial domains and truncated immunogens. In this study, we also observe T cell cross-reactivity to SARS-CoV-1. Further evaluation of these shared responses is likely of interest and may inform development of vaccines targeting related betacoronaviruses. Long-lived SARS-CoV specific T cells have been reported (Le Bert et al., 2020; Yang et al., 2006). Additional studies of SARS-CoV-2 infected patients will provide important insight towards immunity and long-term protection against COVID-19 disease.

In addition to T cells, we demonstrated that INO-4800 induced durable antibody responses that rapidly increased following SARS-CoV-2 challenge. The vaccine induced robust neutralizing antibody responses against both D614 and G614 SARS-CoV-2 variants. While the D/G 614 site is outside the RBD, it has been suggested that this shift has the potential to impact vaccine-elicited antibodies (Korber B et al., 2020) and possibly virus infectivity. Our data shows induction of comparable neutralization titers between D614 and G614 variants and that these responses are similarly recalled following SARS-CoV-2 challenge. We additionally observe a positive relationship between vaccine-induced antibody binding to the RBD and neutralizing antibody titers (**Supplementary Figure 5**, R^2^=0.69).

Overall, the current challenge study provides a snapshot of anamnestic protective efficacy several months post-vaccination with a small cohort of animals. The data support further study of the SARS-CoV-2 DNA vaccine candidate INO-4800 which is currently in clinical trials. The persistence of SARS-CoV-2 immunity following natural infection is unknown and recent studies suggest natural immunity may be short lived (Long et al., 2020). Given that many people exhibit asymptomatic or mild disease and recover without developing significant antibody responses, additional study of vaccines that induce immunological memory for T and B cell responses is clearly of interest. The immune responses and protection induced by simple ID delivery of INO-4800 are promising and provide important information to advancing SARS-CoV-2 vaccine development.

## Acknowledgements

The studies described in this manuscript were funded by a grant from the Coalition for Epidemic Preparedness Innovations (CEPI). The authors would like to additional thank Olivia Bedoya, Gloria Mendez, Francisco Vega Vega and Jon Schantz at Inovio Pharmaceuticals for their assistance, and Jack Greenhouse and Tammy Putmon-Taylor at Bioqual for their expert assistance.

## Author Contributions

Conceptualization, A.P., J.W., J.J.K., S.R., T.R.F.S, D.B.W., K.E.B

Methodology, A.P., J.W., E.L.R., K.S., E.P., E.N.G., D.W.K

Investigation, A.P., J.W., E.L.R., K.S., E.P., E.N.G., I.M., M.P., Z.E., S.N.W., D.G., P.B., A.D., Z.X., D.E., S.M.R., L.P., K.Y.K., A.C., N.C., B.F., E.T-R., A.D., J.C., A.G., N.J.T., V.M.A., Y.D., F.I.Z., D.W.K

Resources, K.M., D.W.K, H.A., M.G.L.

Writing – Original Draft, A.P., J.W., T.R.F.S.

Writing – Review & Editing, A.P., J.W., E.L.R., E.P., E.N.G., M.P., P.B., J.J.K., L.M.H., S.R., T.R.F.S., D.B.W.

Supervision: A.P., J.W., S.R., T.R.F.S., D.B.W., K.E.B

Project Administration: A.P., J.W., J.J.K., L.M.H., S.R., T.R.F.S., D.B.W., K.E.B.

Funding Acquisition: J.J.K., L.M.H., D.B.W., K.E.B.

## Declaration of Interests

**A.P., E.L.R., E.P., E.N.G., M.P., S.N.W., P.B., Z.X., S.M.R., K.Y.K., N.C., E.T-R., J.C., N.J.T., K.M., D.K.W.,** declare no competing interests. **J.W., K.S., I.M., Z.E., D.G., A.D., D.E., A.G., V.M.A., J.J.K., L.M.H., S.R., T.R.F.S., K.E.B.** are employees of Inovio Pharmaceuticals and as such receive salary and benefits, including ownership of stock and stock options, from the company. **D.B.W**. discloses the following paid associations with commercial partners: GeneOne (Consultant), Geneos (Advisory Board), Astrazeneca (Advisory Board, Speaker), Inovio (BOD, SRA, Stock), Pfizer (Speaker), Merck (Speaker), Sanofi (Advisory Board), BBI (Advisory Board).

## Methods

### DNA vaccine, INO-4800

The highly optimized DNA sequence encoding SARS-CoV-2 IgE-spike was created using Inovio’s proprietary in silico Gene Optimization Algorithm to enhance expression and immunogenicity (Smith et al., 2020). The optimized DNA sequence was synthesized, digested with BamHI and XhoI, and cloned into the expression vector pGX0001 under the control of the human cytomegalovirus immediate-early promoter and a bovine growth hormone polyadenylation signal.

### Animals

All rhesus macaque experiments were approved by the Institutional Animal Care and Use Committee at Bioqual (Rockville, Maryland), an Association for Assessment and Accreditation of Laboratory Animal Care (AAALAC) International accredited facility. Blood was collected for blood chemistry, PBMC isolation, and serological analysis. Bronchoalveolar lavage (BAL) was collected on Week 8 to assay lung antibody levels and on Days 1, 2, 4, 7 post challenge to assay lung viral loads.

### Immunizations, sample collection and viral challenge

Ten Chinese rhesus macaques (ranging from 4.55kg-5.55kg) were randomly assigned to be immunized (3 males and 2 females) or naïve (2 males and 3 females). Immunized macaques received two 1 mg injections of SARS-CoV-2 DNA vaccine, INO-4800, at week 0 and 4 by the minimally invasive ID-EP administration using the CELLECTRA 2000^®^ Adaptive Constant Current Electroporation Device with a 3P array (Inovio Pharmaceuticals). Animals were challenged with SARS-Related Coronavirus 2, Isolate USA-WA1/2020 (BEI Resources NR-52281. The following reagent was deposited by the Centers for Disease Control and Prevention and obtained through BEI Resources, NIAID, NIH: SARS-Related Coronavirus 2, Isolate USA-WA1/2020, NR-52281. Blood was collected at indicated time points to analyse blood chemistry and to isolate peripheral blood mononuclear cells (PBMC), and serum. Bronchoalveolar lavage was collected at Week 8 and during challenge period to assay viral load. BAL from naïve animals was run as control. At week 17, all animals were challenged with 1.2×10^8^ VP (1.1×10^4^ PFU) SARS-CoV-2. Virus was administered as 1 ml by the intranasal route (0.5 ml in each nostril) and 1 ml by the intratracheal route. Nasal swabs were collected during the challenge period using Copan flocked swabs and placed into 1 ml PBS.

### Peripheral blood mononuclear cell isolation

Blood was collected from each macaque into sodium citrate cell preparation tubes (CPT, BD Biosciences). The tubes were centrifuged to separate plasma and lymphocytes, according to the manufacturer’s protocol. Samples were transported by same-day shipment on cold-packs from Bioqual to The Wistar Institute for PBMC isolation. PBMCs were washed and residual red blood cells were removed using ammonium-chloride-potassium (ACK) lysis buffer. Cells were counted using a ViCell counter (Beckman Coulter) and resuspended in RPMI 1640 (Corning), supplemented with 10% fetal bovine serum (Atlas), and 1% penicillin/streptomycin (Gibco). Fresh cells were then plated for IFNγ ELISpot Assays and flow cytometry.

### IFN-γ Enzyme-linked immunospot (ELISpot)

Monkey interferon gamma (IFN-γ) ELISpot assay was performed to detect cellular responses. Monkey IFN-γ ELISpotPro (alkaline phosphatase) plates (Mabtech, Sweeden, Cat#3421M-2APW-10) were blocked for a minimum of 2 hours with RPMI 1640 (Corning), supplemented with 10% FBS and 1% pen/strep (R10). Following PBMC isolation, 200 000 cells were added to each well in the presence of 1) peptide pools (15-mers with 9-mer overlaps) corresponding to the SARS-CoV-1, SARS-CoV-2, or MERS-CoV spike proteins (5 μg/mL/well final concentration), 2) R10 with DMSO (negative control), or 3) anti-CD3 positive control (Mabtech, 1:1000 dilution). All samples were plated in triplicate. Plates were incubated overnight at 37°C, 5% CO2. After 18-20 hours, the plates were washed in PBS and spots were developed according to the manufacturer’s protocol. Spots were imaged using a CTL Immunospot plate reader and antigen-specific responses were determined by subtracting the number of spots in the R10+DMSO negative control well from the wells stimulated with peptide pools.

### Antigen Binding ELISA

Serum and BAL was collected at each time point was evaluated for binding titers. Ninety-six well immunosorbent plates (NUNC) were coated with 1ug/mL recombinant SARS-CoV-2 S1+S2 ECD protein (Sino Biological 40589-V08B1), S1 protein (Sino Biological 40591-V08H), S2 protein (Sino Biological 40590-V08B), or receptor-binding domain (RBD) protein (Sino Biological 40595-V05H) in PBS overnight at 4°C. ELISA plates were also coated with 1 μg/mL recombinant SARS-CoV S1 protein (Sino Biological 40150-V08B1) and RBD protein (Sino Biological 40592-V08B) or MERS-CoV spike (Sino Biological 40069-V08B). Plates were washed four times with PBS + 0.05% Tween20 (PBS-T) and blocked with 5% skim milk in PBS-T (5% SM) for 90 minutes at 37°C. Sera or BAL from INO-4800 vaccinated and control macaques were serially diluted in 5% SM, added to the washed ELISA plates, and then incubated for 1 hour at 37°C. Following incubation, plates were washed 4 times with PBS-T and an anti-monkey IgG conjugated to horseradish peroxidase (Southern Biotech 4700-5) 1 hour at 37°C. Plates were washed 4 times with PBS-T and one-step TMB solution (Sigma) was added to the plates. The reaction was stopped with an equal volume of 2N sulfuric acid. Plates were read at 450nm and 570nm within 30 minutes of development using a Biotek Synergy2 plate reader.

### ACE2 Competition ELISA-Non-human primates

96-well half area plates (Corning) were coated at room temperature for 3 hours with 1 μg/mL PolyRab anti-His antibody (ThermoFisher, PA1-983B), followed by overnight blocking with blocking buffer containing 1x PBS, 5% SM, 1% FBS, and 0.2% Tween-20. The plates were then incubated with 10 μg/mL of His6x-tagged SARS-CoV-2, S1+S2 ECD (Sinobiological, 40589-V08B1) at room temperature for 1-2 hours. NHP sera (Day 0 or Week 6) was serially diluted 3-fold with 1XPBS containing 1% FBS and 0.2% Tween and pre-mixed with huACE2-IgMu at constant concentration of 0.4 μg/ml. The pre-mixture was then added to the plate and incubated at room temperature for 1-2 hours. The plates were further incubated at room temperature for 1 hour with goat anti-mouse IgG H+L HRP (A90-116P, Bethyl Laboratories) at 1:20,000 dilution followed by addition of one-step TMB ultra substrate (ThermoFisher) and then quenched with 1M H2SO4. Absorbance at 450nm and 570nm were recorded with BioTEK plate reader.

### Flow cytometry-based ACE2 receptor binding inhibition assay

HEK-293T cells stably expressing ACE2-GFP were generated using retroviral transduction. Following transduction, the cells were flow sorted based on GFP expression to isolate GFP positive cells. Single cell cloning was done on these cells to generate cell lines with equivalent expression of ACE2-GFP. To detect inhibition of spike binding to ACE2, 2.5 μg/ml S1+S2 ECD-his tagged (Sino Biological, Catalog #40589-V08B1) was incubated with serum collected from vaccinated animals at indicated time points and dilutions on ice for 60 min. This mixture was then transferred to 150,000 293T-ACE2-GFP cells and incubated on ice for 90 min. Following this, the cells were washed 2x with PBS followed by staining for surelight^®^ APC conjugated anti-his antibody (Abcam, ab72579) for 30 min on ice. As a positive control, spike protein was pre-incubated with recombinant human ACE2 before transferring to 293T-ACE2-GFP cells. Data was acquired using a BD LSRII and analyzed by FlowJo (version 10).

#### Pseudovirus Neutralization Assay

SARS-CoV-2 pseudovirus was produced using HEK293T cells transfected with GeneJammer (Agilent) using IgE-SARS-CoV-2 spike plasmid (Genscript) and pNL4-3.Luc.R-E-plasmid (NIH AIDS reagent) at a 1:1 ratio. Forty-eight hours post transfection, supernatant was collected, enriched with FBS to 12% final volume, steri-filtered (Millipore Sigma), and aliquoted for storage at −80°C. SARS-Cov-2 pseudovirus neutralization assay was set up using D10 media (DMEM supplemented with 10%FBS and 1X Penicillin-Streptomycin) in a 96 well format. CHO cells stably expressing ACE2 were used as target cells (Creative Biolabs, Catalog No. VCeL-Wyb019). SARS-Cov-2 pseudovirus were titered to yield greater than 20 times the cells only control relative luminescence units (RLU) after 72h of infection. 10,000 CHO-ACE2 cells/well were plated in 96-well plates in 100ul D10 media and rested overnight at 37°C and 5% CO2 for 24 hours. The following day, sera from INO-4800 vaccinated and control groups were heat inactivated and serially diluted. Sera were incubated with a fixed amount of SARS-Cov-2 pseudovirus for 90 minutes at RT. The sera+virus mix was then added to the plated CHO-ACE2 cells and allowed to incubate in a standard incubator (37% humidity, 5% CO2) for 72h. Cells were then lysed using Britelite plus luminescence reporter gene assay system (Perkin Elmer Catalog no. 6066769) and RLU were measured using the Biotek plate reader. Neutralization titers (ID50) were calculated using GraphPad Prism 8 and defined as the reciprocal serum dilution at which RLU were reduced by 50% compared to RLU in virus control wells after subtraction of background RLU in cell control wells.

### Viral RNA assay

RT-PCR assays were utilized to monitor viral loads, essentially as previously described (Abnink P et al 2019 Science). Briefly, RNA was extracted using a QIAcube HT (Qiagen,Germany) and the Cador pathogen HT kit from bronchoalveolar lavage (BAL) supernatant and nasal swabs. RNA was reverse transcribed using superscript VILO (Invitrogen) and ran in duplicate using the QuantStudio 6 and 7 Flex Real-Time PCR System (Applied Biosystems) according to manufacturer’s specifications. Viral loads were calculated of viral RNA copies per mL or per swab and the assay sensitivity was 50 copies. The target for amplification was the SARS-CoV2 N (nucleocapsid) gene. The primers and probes for the targets were:

2019-nCoV_N1-F: 5’-GACCCCAAAATCAGCGAAAT-3’
2019-nCoV_N1-R: 5’-TCTGGTTACTGCCAGTTGAATCTG-3’
2019-nCoV_N1-P: 5’-FAM-ACCCCGCATTACGTTTGGTGGACC-BHQ1-3’

### Subgenomic mRNA assay

SARS-CoV-2 E gene subgenomic mRNA (sgmRNA) was assessed by RT-PCR using an approach similar to previously described (Wolfel R et al. Nature 2020). To generate a standard curve, the SARS-CoV-2 E gene sgmRNA was cloned into a pcDNA3.1 expression plasmid; this insert was transcribed using an AmpliCap-Max T7 High Yield Message Maker Kit (Cellscript) to obtain RNA for standards. Prior to RT-PCR, samples collected from challenged animals or standards were reverse-transcribed using Superscript III VILO (Invitrogen) according to the manufacturer’s instructions. A Taqman custom gene expression assay (ThermoFisher Scientific) was designed using the sequences targeting the E gene sgmRNA (18). Reactions were carried out on a QuantStudio 6 and 7 Flex Real-Time PCR System (Applied Biosystems) according to the manufacturer’s specifications. Standard curves were used to calculate sgmRNA in copies per ml or per swab; the quantitative assay sensitivity was 50 copies per ml or per swab.

## Notes

### Competing Interest Statement

A.P., E.R., E.P., E.N.G., M.P., S.N.W., P.B., Z.X., S.R., K.Y.K., N.C., E.T-R., J.C., N.J.T., K.M., D.K.W., declare no competing interests. J.W., K.S., I.M., Z.E., D.G., A.D., D.E., A.G., V.M.A., J.J.K., L.M.H., S.R., T.R.F.S., K.E.B. are employees of Inovio Pharmaceuticals and as such receive salary and benefits, including ownership of stock and stock options, from the company. D.B.W. discloses the following paid associations with commercial partners: GeneOne (Consultant), Geneos (Advisory Board), Astrazeneca (Advisory Board, Speaker), Inovio (BOD, SRA, Stock), Pfizer (Speaker), Merck (Speaker), Sanofi (Advisory Board), BBI (Advisory Board).

