## Supplemental figures for "Intradermal-delivered DNA vaccine provides anamnestic protection in a rhesus macaque SARS-CoV-2 challenge model"

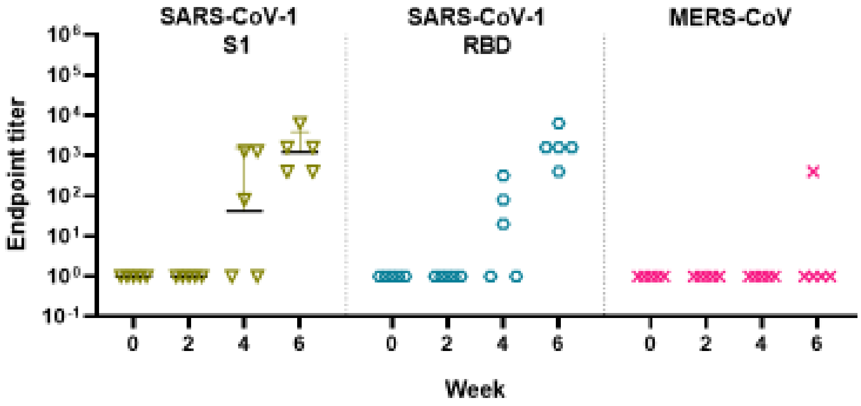


**Supplemental Figure 1. Serum IgG cross-reactivity to SARS-CoV and MERS-CoV spike protein.** IgG binding was measured in sera from INO-4800 vaccinated rhesus macaques to SARS-CoV S1 and RBD and MERS-CoV S1 protein antigen.

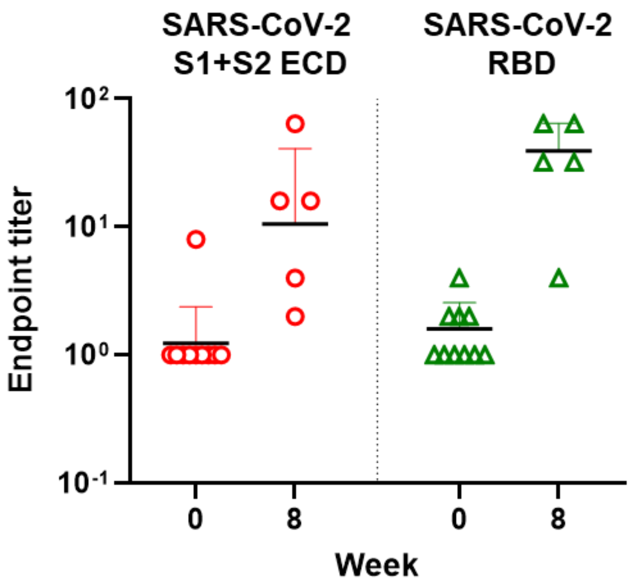


**Supplemental Figure 2**. **Bronchoalveolar lavage (BAL) IgG reactive to SARS-CoV-2 S protein antigens.** BAL samples collected from vaccinated animals were assessed for SARS-CoV-2 reactive IgG binding to the full length SARS-CoV-2 spike protein and the RBD domain.

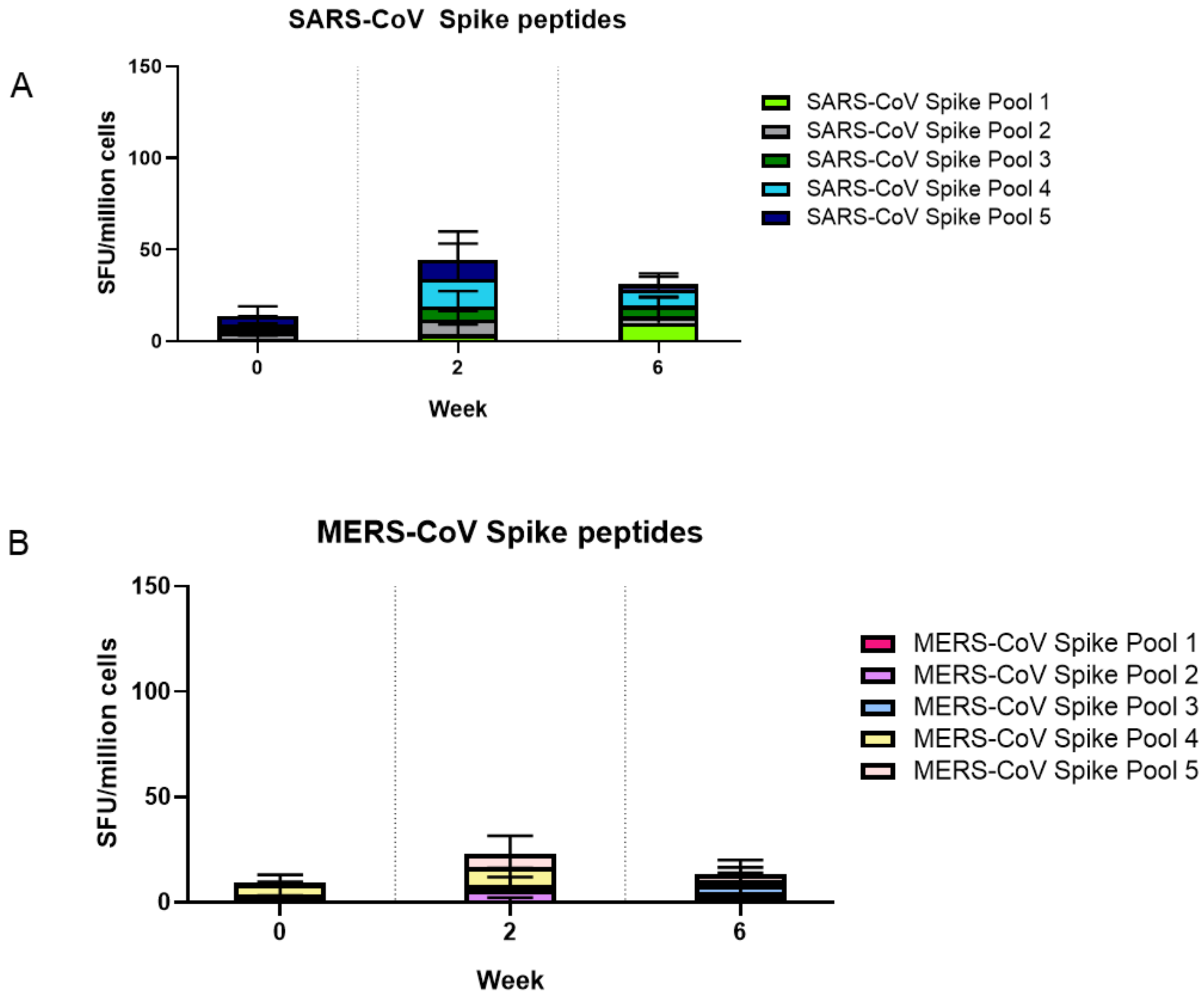


**Supplemental Figure 3. Cellular response cross-reactivity to SARS-CoV and MERS-CoV spike protein.** PBMC responses were analyzed by IFNγ ELISpot after stimulation with overlapping peptide pools spanning the SARS-CoV-1 spike protein (**A**) and MERS-CoV spike protein (**B**). Bars represent the mean + SD.

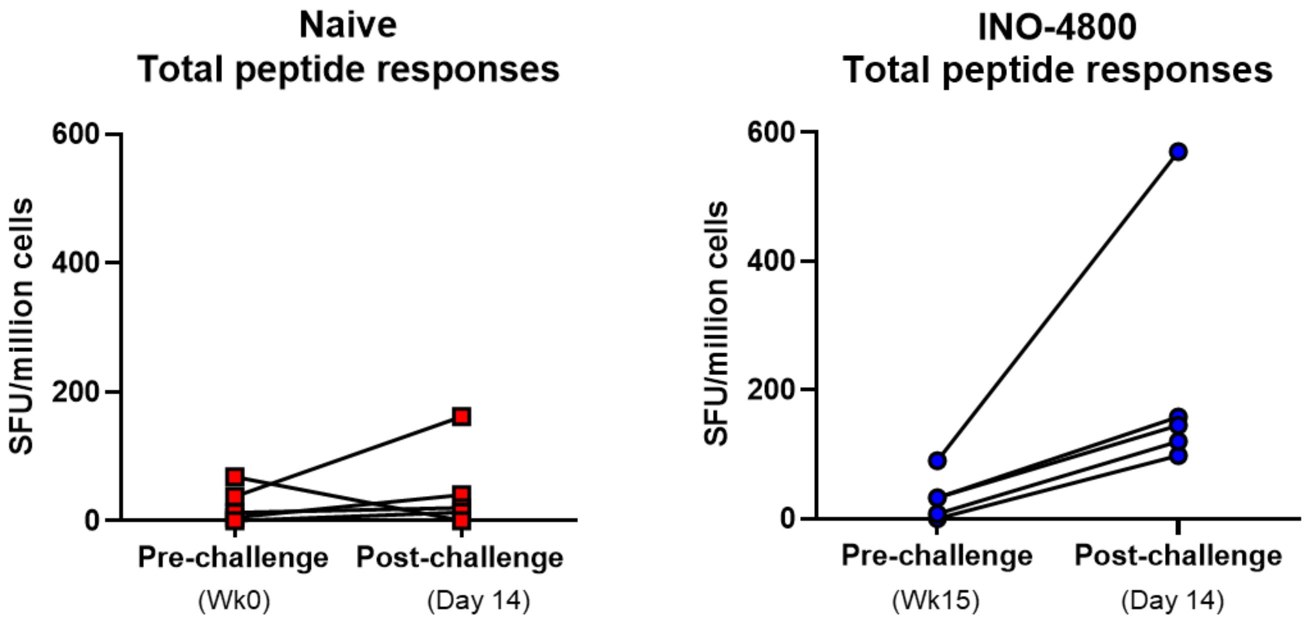


**Supplemental Figure 4. Recall of cellular immune responses after viral challenge in individual rhesus macaques.** Cellular responses were analyzed pre and post viral challenge by IFNγ ELISpot in PBMCs stimulated with overlapping peptide pools spanning the SARS-CoV-2 spike protein. Left panel naïve animals, right panel INO-4800 vaccinated animals

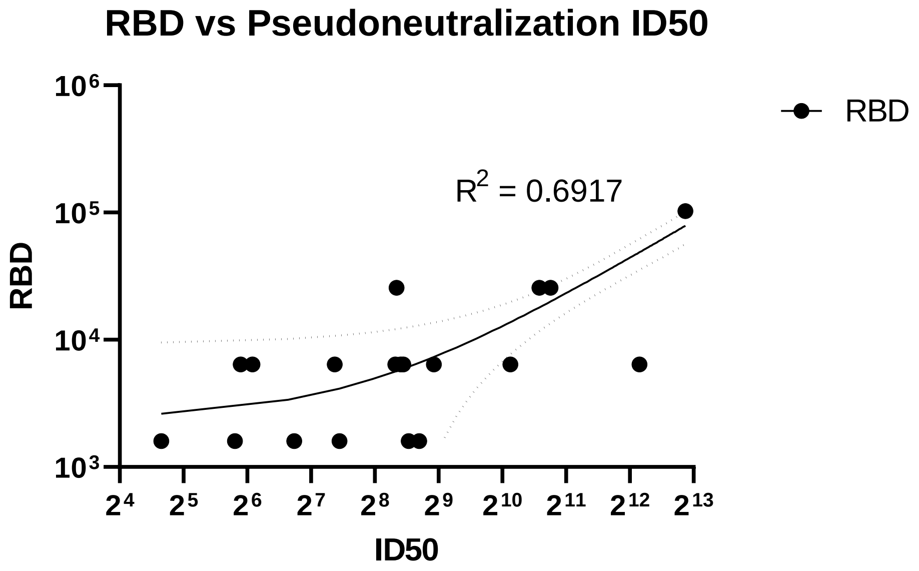


**Supplemental Figure 5. Simple linear regression analysis comparing total IgG antibodies against the RBD and neutralizing antibodies raised in DNA-vaccinated macaques.**
